# Tumor-specific antibodies elicited by engineered bacteria promote bladder cancer immunotherapy

**DOI:** 10.1101/2024.10.24.620122

**Authors:** Mathieu Rouanne, Noah Chen, Dylan L. Mariuzza, Fangda Li, Kenia de los Santos-Alexis, Thomas M. Savage, Rosa L. Vincent, Cathy L. Mendelsohn, Tal Danino, Nicholas Arpaia

## Abstract

The intratumoral microbiome has recently emerged as a new hallmark of cancer, with implications for response or resistance to therapy. While bacteria can either promote or inhibit cancer growth, intratumoral bacteria can also be engineered using synthetic biology to remodel the tumor microenvironment. Here, we engineered the probiotic bacterium *E. coli* Nissle 1917 (EcN) to express the human chemokine CXCL13, a critical component of germinal center (GC) formation. The GC reaction is a fundamental aspect of adaptive immunity by which antibody affinity develops in secondary lymphoid organs for defense against pathogens. Using orthotopic models of bladder cancer, engineered CXCL13-expressing EcN colonized bladder tumors and elicited GC responses in bladder tumor–draining lymph nodes after intravesical delivery. Furthermore, when combined with PD-1 blockade, engineered EcN amplified the antitumor antibody response and promoted long-term survival and protective immunity upon tumor rechallenge. Thus, we demonstrate that synthetically engineered CXCL13-expressing EcN can enhance the efficacy of PD-1 checkpoint blockade immunotherapy by amplifying tumor-specific humoral immunity.

## Main text

Recent evidence suggests that anti-programmed cell death protein 1 (PD-1) immune checkpoint blockade (ICB), a therapeutic monoclonal antibody which promotes durable tumor regression across multiple cancer types, acts not only at the tumor site but also in the draining lymph nodes, where it activates a germinal center (GC) response (*1–3*). The GC reaction is a fundamental component of adaptive immunity, during which antibody affinity and the generation of immunological memory occur in lymphoid organs in response to local antigenic challenge (*4*). Although intratumoral microbes have been identified as an intrinsic component of the tumor microenvironment (TME) across human cancer types, their effect on the host humoral immune response during ICB immunotherapy remains poorly understood, which further hinders the ability to develop more effective therapeutic interventions (*5–7*).

Bacillus Calmette–Guérin (BCG) immunotherapy for bladder cancer is the only bacterial cancer therapy approved for clinical use. Upon intravesical delivery, BCG induces an antitumor effect that is dependent on tumor-specific CD4^+^ T cells, similarly to most recently employed ICB strategies with regard to the induction of tumor-specific immunity (*8, 9*). Emerging evidence has revealed that the presence of intratumoral *E. coli* in human tumors (including bladder cancer) was associated with improved survival in response to PD-1 blockade in independent clinical cohorts, suggesting a link between intratumoral bacteria and response to immunotherapy (*10, 11*). Indeed, *E. coli* is the most common bacteria observed in the urinary system, and has been found to induce robust immunogenic cellular and humoral immune responses during urinary tract infections (*12, 13*). Here, we engineer a probiotic, non-pathogenic strain of *E. coli* Nissle 1917 (EcN) to express the human chemokine CXCL13 (EcN^hCXCL13^) for intravesical delivery. The CXCL13–CXCR5 chemokine axis plays a central role in GC formation by organizing B cell follicles, and recently has been associated with immunotherapy response across multiple tumor types, including bladder cancer (*1, 14–17*). In addition, circulating CXCL13 levels, a plasma biomarker of GC activity, has been found to be positively correlated with early response to PD-1 blockade in bladder cancer patients (*10, 18*).

Here, we show that intravesically delivered EcN^hCXCL13^ colonize bladder tumors and promote GC responses that augment the production of antitumor IgG antibodies upon combination with PD-1 blockade, which in turn enhances ICB therapeutic efficacy. Thus, probiotics synthetically engineered to express immunotherapeutic payloads in combination with PD-1 blockade represents a promising strategy to leverage immune activation for the treatment of bladder cancer.

## Results

### Engineering E. coli Nissle 1917 to release the human chemokine CXCL13 and promote the GC reaction in bladder tumor–draining lymph nodes

To effectively deliver the human chemokine CXCL13 (hCXCL13) to the bladder TME, we engineered a well-characterized probiotic strain, *Escherichia coli* Nissle 1917 (EcN), equipped with a previously-described synchronized lysis integrated circuit (SLIC) for quorum sensing–triggered release of plasmid-encoded hCXCL13, EcN^hCXCL13^ **(Fig.1A and fig.S1A)** (*19–22*). Specifically, bacteria harboring this integrated circuit grow until they reach a critical population density within the TME, therefore triggering cyclic lysis events that release co-encoded payloads, such as chemokines, in situ. First, we confirmed that release of hCXCL13 was SLIC dependent and EcN^hCXCL13^ demonstrated no difference in growth rate relative to EcN^WT^ in non-lysing conditions **(fig. S1C, and B)**. Taken together, these data suggest that incorporation of both a hCXCL13-expressing plasmid and the integrated SLIC system did not affect bacterial growth as compared to the wild-type EcN. To further evaluate the functionality of hCXCL13 produced by the engineered bacteria, we performed a chemotaxis assay, in which the migration of human B cells expressing high levels of the chemokine receptor CXCR5 was evaluated in response to lysates of EcN^hCXCL13^, EcN expressing mouse CXCL13 (EcN^mCXCL13^), or control EcN (*23*). Compared to control EcN lysate, human B cells significantly migrated in response to lysates of both EcN^hCXCL13^ and EcN^mCXCL13^ **(fig.S1D)**. Similarly, mouse splenocytes migrated to a greater extent to lysate from EcN^hCXCL13^ **(fig.S1D)**. Thus, we used EcN^hCXCL13^ to facilitate future translation to human studies.

**Figure 1.**
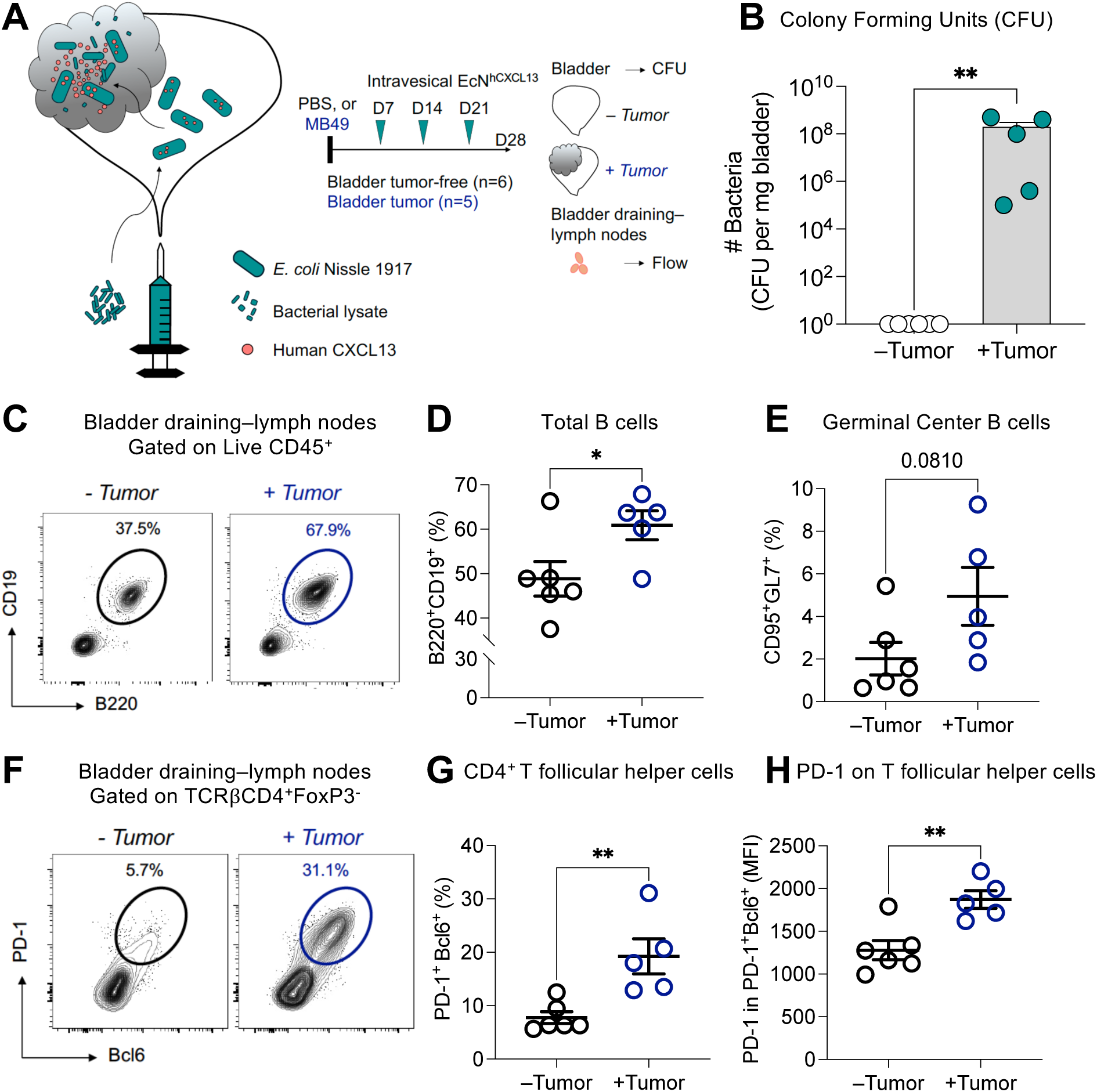
Engineering *E. coli* Nissle 1917 to release the human chemokine CXCL13 and promote germinal center reaction in bladder tumor–draining lymph nodes. (**A**) Schematic demonstrating the release of the human chemokine CXCL13 in bladder tumors after tumor colonization by engineered probiotic *E.coli* Nissle 1917 upon intravesical delivery (left panel). PBS or MB49 cells (2 x 10^5^) were implanted in the bladder of C57/BL6 mice. Intravesical delivery of 3 x 10^6^ CFU of EcN^hCXCL13^ was performed on days 7, 14, and 21. On day 28, supernatant from freshly dissociated tumor-free (n=6) or tumor-bearing (n=5) bladders was plated on LB agar plates containing kanamycin. Colony-forming units (CFU) were counted after overnight culture at 37°C and computed as CFU per mg of bladder tissue. Bladder tumor–draining lymph nodes were dissociated for immune phenotyping by flow cytometry (right panel). (**B**) Quantification of the total number of CFU per bladder weight (mg) after overnight culture at 37°C on LB agar plate with kanamycin. Data shown represent two independent experiments with a total of 5-6 mice per group (**P < 0.01, two-tailed unpaired Mann Whitney test). (**C**) Representative dot plots showing total CD19^+^B220^+^ B cells in draining lymph nodes from bladder tumor-free (left panel) and bladder tumor-bearing (right panel) mice. (**D**) Flow cytometric quantification of total CD19^+^B220^+^ B cells among live CD45^+^ cells in draining lymph nodes from bladder-tumor-free and bladder-tumor bearing mice (*P < 0.05, two-tailed unpaired Student’s *t* test). (**E**) Flow cytometric quantification of CD95^+^GL7^+^ germinal center B cells among total B cells in draining lymph nodes from bladder tumor-free and bladder tumor-bearing mice (two-tailed unpaired Student’s *t* test). (**F**) Representative dot plots showing Bcl6^+^PD-1^+^ T follicular helper cells among TCRβ^+^CD4^+^FoxP3^-^ cells in draining lymph nodes from bladder tumor-free (left panel) and bladder tumor-bearing (right panel) mice. (**G**) Flow cytometric quantification of Bcl6^+^PD-1^+^ T follicular helper cells among TCRβ^+^CD4^+^FoxP3^-^ cells in draining lymph nodes from bladder tumor-free and bladder tumor-bearing mice (**P < 0.01, two-tailed unpaired Student’s *t* test). (**H**) Mean fluorescence intensity (MFI) quantification of PD-1 molecules expressed at the surface of Bcl6^+^PD-1^+^ T follicular helper cells by flow cytometry in draining lymph nodes from bladder tumor-free and bladder tumor-bearing mice (**P < 0.01, two-tailed unpaired Student’s *t* test).

Encouraged by our observations in vitro, we next sought to interrogate the biological activity of EcN^hCXCL13^ in vivo using orthotopic models of bladder cancer. To generate bladder tumors, we intravesically implanted MB49 cells in syngeneic wild-type mice, a well-studied orthotopic mouse model of bladder cancer with an aggressive phenotype (*24, 25*). To assess the ability of EcN^hCXCL13^ to effectively and specifically colonize bladder tumors upon intravesical delivery, bladder tumor–bearing and tumor-free mice were exposed to repeated intravesical instillations of engineered bacteria once a week for 3 weeks **(Fig.1A)** Notably, colonies of EcN^hCXCL13^ were observed only in bladder tumors and were not found in tumor-free bladders after intravesical delivery **(Fig.1B)**. Furthermore, bacterial colonies isolated from freshly dissociated bladder tumors were able to grow under selective antibiotic pressure and release the human chemokine CXCL13, confirming their ability to maintain therapeutic plasmids while growing in vivo **(fig. S1E)**. These observations are consistent with our hypothesis that bacteria preferentially colonize bladder tumors while sparing non-tumoral tissue and consistent with prior preclinical data (*19–22*). Furthermore, we monitored body weight variation of bladder tumor–bearing mice from the start of bacteria treatment and observed no significant weight loss in mice treated with EcN^hCXCL13^, EcN^−^ control (empty vector), or PBS (**fig. S1F**), confirming the favorable safety profile of our therapeutic approach.

To characterize the immune response to our therapeutic strain, we performed immunophenotyping of T cell and B cell infiltrates in the draining lymph nodes from bladder tumor-bearing and tumor-free mice after intravesical delivery of EcN^hCXCL13^, with the aim of assessing the GC response. We detected an increased frequency of total B cells and GC B cells, as well as an increase in Tfh cells, in bladder tumor–draining lymph nodes as compared with bladder tumor-free mice **(Fig.1, C to G)**. Notably, we found a significant increase of cell surface PD-1 molecules on Tfh cells in bladder tumor–draining lymph nodes. These data suggest that intravesical delivery of EcN^hCXCL13^ actively promotes the GC reaction in draining lymph nodes from bladder tumor–bearing mice, most likely due to the in situ release of hCXCL13 in bladder tumors colonized by EcN^hCXCL13^ **(Fig.1H)**. Together, these observations demonstrate both the safety and ability for intravesically-delivered EcN^hCXCL13^ to elicit the GC reaction in orthotopic models of bladder cancer.

### EcN^hCXCL13^ synergizes with PD-1 blockade to promote long-term survival and durable protection in an orthotopic model of advanced bladder cancer

Next, we hypothesized that EcN^hCXCL13^ could sensitize tumors to PD-1 blockade by activating the GC reaction in bladder tumor–draining lymph nodes in response to PD-1 binding on Tfh cells. Of note, in the absence of PD-1 blockade the median survival of MB49 bladder tumor–bearing mice treated with intravesical delivery of EcN^hCXCL13^ at days 3, 6, and 9 post-tumor implantation was 35 days, compared with 31 and 27 days for mice treated with EcN^−^ or PBS, respectively **(Fig.2, A and B)**. Although we did not observe significant antitumor activity of EcN^hCXCL13^ as a single agent in the MB49 orthotopic model, in a separate established orthotopic model of bladder cancer using the UPPL1541 cell line, as compared with animals treated with control EcN or PBS, we consistently noted a spatial reorganization of immune cells in the bladder mucosa, with the induction of immune aggregates of CD4^+^ T cells, CD138^+^ plasma cells and F4/80^+^ macrophages 4 weeks after repeated intravesical instillations of EcN^hCXCL13^ **(fig. S2, A and B)**(*26*). These data suggest that EcN^hCXCL13^ was biologically and immunologically active at the mucosal surface of the bladder upon intravesical delivery but was not sufficient as a single therapeutic agent to induce a significant survival benefit.

**Figure 2.**
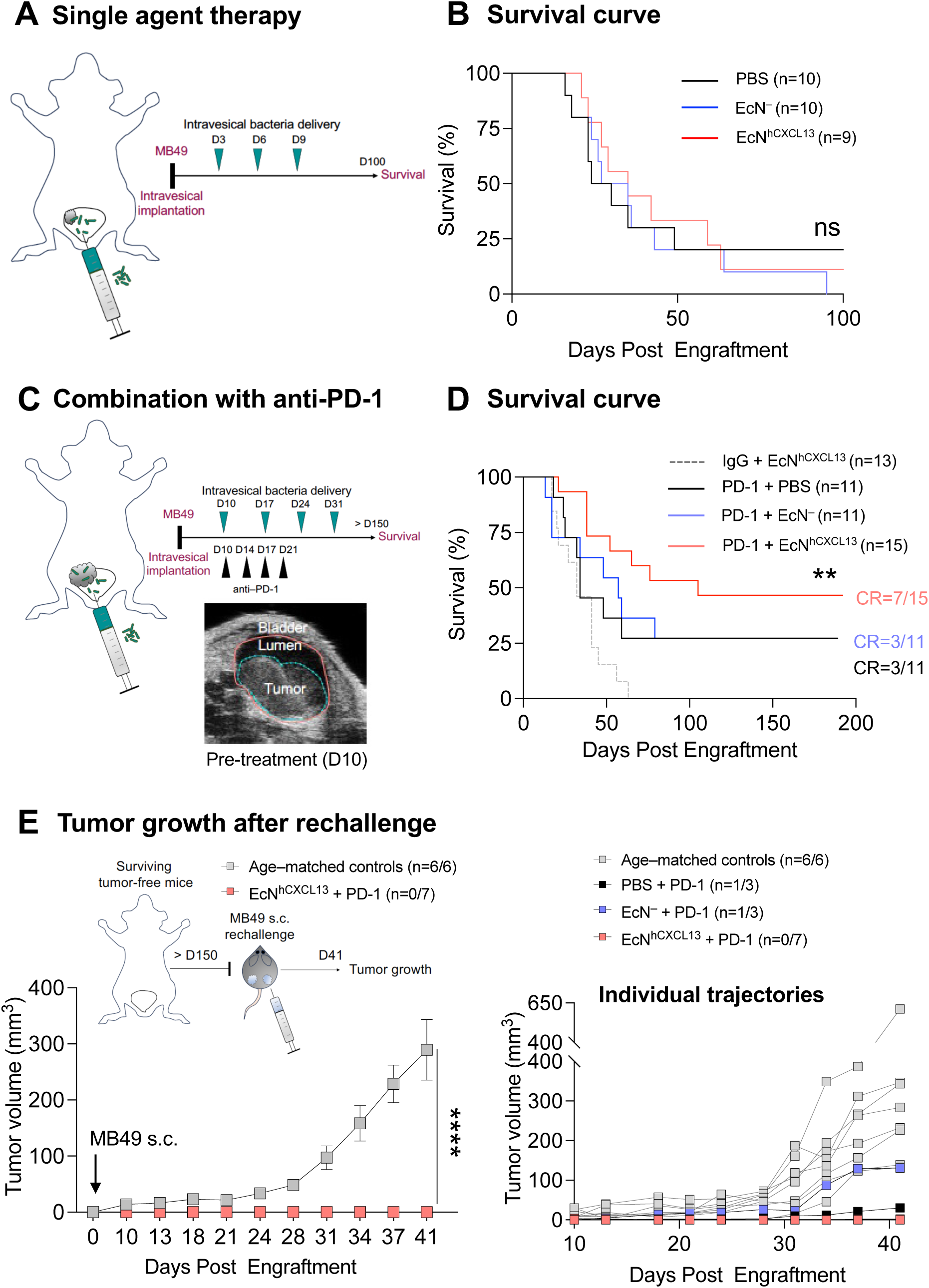
EcN^hCXCL13^ synergizes with PD-1 blockade to promote long-term survival and durable protection in an orthotopic model of advanced bladder cancer. (**A**) MB49 cells (2 x 10^5^) were implanted in the bladder of wild-type C57BL/6 female mice. Intravesical delivery of either 3 x 10^6^ CFU of EcN^hCXCL13^, EcN^−^ control (empty vector), or PBS was performed at days 3, 6, and 9. Survival was monitored until day 100. (**B**) Survival curves are shown for each treatment group. Data represent one experiment with n=9-10 mice per group (log-rank test). (**C**) MB49 cells (2 x 10^5^) were implanted in the bladder of wild-type C57BL/6 female mice. At day 10, the volume of bladder tumors was monitored by ultrasound imaging. Mean pre-treatment tumor volume was ∼30 mm^3^. Intravesical delivery of either 3 x 10^6^ CFU of EcN_slic_^hCXCL13^, EcN^−^ control (empty vector), or PBS was performed at day 10, 17, 24, 31. Intraperitoneal injections of PD-1 blockade were done on days 10, 14, 17, and 21. Isotype control IgG was combined with EcN^hCXCL13^ as an additional control. Survival was monitored for > 150 days. (**D**) Survival curves are shown for each treatment group. Tumor clearance was confirmed by ultrasound imaging of the bladder in all surviving mice > 150 days. Data represent three independent experiments with n=11-15 mice per group (log-rank test). (**E**) To assess long-term antitumor control, surviving mice from (**D**) or age-matched control mice were subcutaneously implanted into both hind flanks with MB49 cells (2 x 10^5^). Tumor volume was monitored for 41 days post-engraftment (top left panel). Mean tumor growth trajectories are shown in surviving mice previously treated with EcN^hCXCL13^ plus PD-1 blockade and age-matched controls (bottom left panel) (****P < 0.0001, two-way ANOVA with Holm-Sidak post hoc test). Individual trajectories of tumor growth are shown in mice from all treatment groups and age-matched controls. MB49 engraftment was observed in mice previously treated with PBS plus PD-1 blockade (n=1/3) and EcN^−^ plus PD-1 blockade (n=1/3). No tumor engraftment was observed in surviving mice previously treated with EcN_slic_^hCXCL13^ plus PD-1 blockade (n=0/7) (right panel).

To further assess the antitumor efficacy of EcN^hCXCL13^ combined with PD-1 blockade in an orthotopic model of advanced bladder cancer with previously described primary resistance to PD-1 blockade, we implanted UPPL1541 cells in the bladder submucosa of mice using an ultrasound–guided approach **(fig. S2C)** (*26, 27*). In this setting, mice were intravesically treated EcN^hCXCL13^, EcN^−^, or PBS at days 7, 10, and 13, along with repeated injections of PD-1 blockade at days 13, 16, and 19. The tumor volumes between pre-treatment day 7 and post-treatment day 21 was evaluated by ultrasound imaging of the bladder to assess the dynamics of bladder tumor growth **(fig. S2C)**. Notably, EcN^hCXCL13^ plus PD-1 blockade significantly restrained tumor growth when compared with UPPL1541 bladder tumors treated with EcN^−^ control or PBS plus PD-1 blockade **(fig. S2D)**. Furthermore, we observed a significant increase in the frequencies of total B cells and CD4^+^ T cells infiltrating the bladder tumors upon EcN^hCXCL13^ plus PD-1 blockade exposure **(fig. S2, E and F)**. Overall, these data support the hypothesis that intravesical delivery of EcN^hCXCL13^ promotes immunotherapy responsiveness in a model of PD-1 resistant bladder tumors.

Next, we aimed to validate the antitumor activity of this combination regimen in the MB49 orthotopic model, by starting the treatment 10 days after bladder implantation of the MB49 cells to generate advanced bladder tumors **(Fig. 2C)**. Intravesical delivery of EcN^hCXCL13^ once a week for 4 weeks was combined with 4 injections of PD-1 blockade every 3–4 days after treatment initiation. We found that EcN^hCXCL13^ and PD-1 blockade significantly extended survival, with ∼47% (n=7/15) of mice surviving and remaining tumor-free after >150 days of follow-up compared to ∼27% (n=3/11) of mice surviving and tumor-free after treatment with EcN^−^ or PBS combined with PD-1 blockade **(Fig. 2D)**. Notably, this correlated with a significant increase in median survival of 105 days after EcN^hCXCL13^ plus PD-1 blockade combination therapy, as compared to 34 and 57 days after EcN^−^ or PBS plus PD-1 blockade, respectively.

To evaluate the robustness and durability of this effect, we rechallenged the surviving mice by subcutaneously injecting MB49 cells on dual flanks as a model of distant metastasis. In contrast to age-matched naïve controls (n=6/6) and surviving mice treated with anti-PD-1 plus EcN^−^ (n=1/3) or PBS (n=1/3), we did not observe tumor engraftment in any (n=0/7) of the surviving mice that had previously been treated with EcN^hCXCL13^ and PD-1 blockade **(Fig. 2E)**. Together, these observations highlight the potent antitumor activity of EcN^hCXCL13^ combined with PD-1 blockade, which significantly enhanced long-term survival and provided durable protection against metastatic rechallenge in an aggressive orthotopic model of bladder cancer.

### T follicular helper cells drive the therapeutic efficacy of EcN^hCXCL13^ plus PD-1 blockade

To gain insights into the mechanisms by which EcN^hCXCL13^ promotes responsiveness to PD-1 blockade, we examined the expression of key molecules associated with the follicular program and function of CD4^+^ Tfh cells. To this end, we first evaluated the proliferation of CD4^+^ T helper cells in bladder tumor–draining lymph nodes in mice receiving intravesical administrations of EcN^hCXCL13^, EcN^−^, PBS, or bacillus Calmette-Guerin (BCG), a live attenuated vaccine approved for bladder cancer therapy which is known to induce CD4^+^ T cell–dependent tumor-specific immunity in an MB49 orthotopic model of bladder cancer **(Fig.3A)** (*8*). Our analysis showed a significant increase in the frequencies of Ki-67^+^CD4^+^FoxP3^−^ T cells after intravesical delivery of EcN^hCXCL13^ as compared with EcN^−^, BCG, or PBS, suggesting that EcN^hCXCL13^ is sufficient to activate naïve CD4^+^ helper T cells in bladder tumor–draining lymph nodes **(Fig.3B)**.

**Figure 3.**
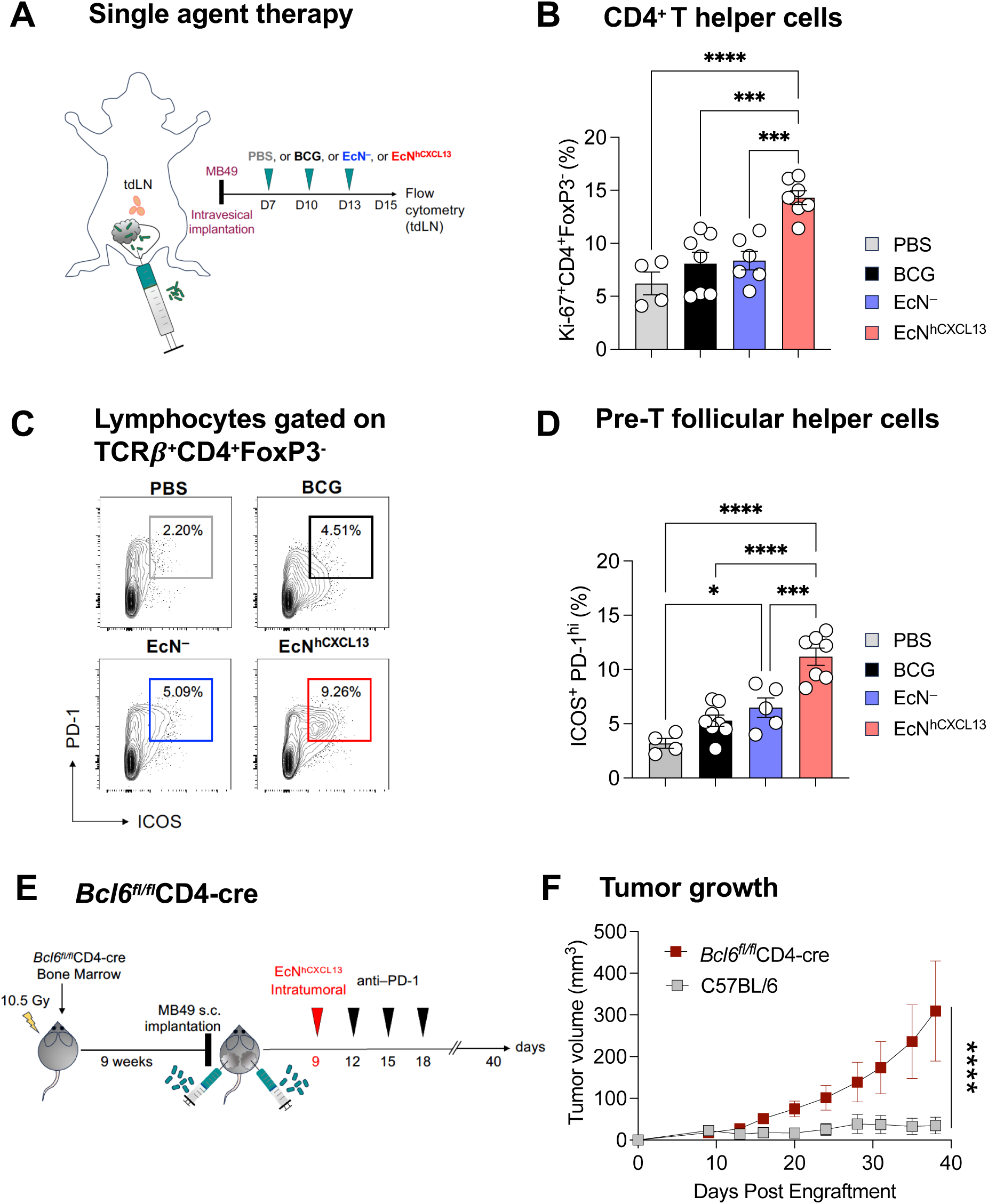
T follicular helper cells drive the therapeutic efficacy of EcN^hCXCL13^ plus PD-1 blockade. (**A**) MB49 cells (2 x 10^5^) were implanted in the bladder of wild-type C57BL/6 female mice. Intravesical delivery of either 3 x 10^6^ CFU of EcN^hCXCL13^, EcN^−^ control (empty vector), BCG, or PBS was performed at day 7, 10, and 13. Bladder tumor–draining lymph nodes were harvested and dissociated for immune phenotyping by flow cytometry 48 hours after the last intravesical delivery. (**B**) Quantification of proliferative effector Ki-67^+^CD4^+^FoxP3^-^ T helper cells in bladder tumor–draining lymph nodes. Data shown represent two independent experiments with a total of 4-7 mice per group. (\*\*\*\**P* < 0.0001, \*\*\**P* < 0.001, two-way ANOVA with Holm-Sidak post hoc test). (**C**) Representative flow plots of CD4^+^FoxP3^-^ T helper cells co-expressing ICOS^+^PD-1^+^ molecules from bladder tumor–draining lymph nodes of mice treated as in (**A**). (**D**) Quantification of activated CD4^+^FoxP3^-^ T helper cells co-expressing ICOS^+^ and PD-1^+^ molecules from bladder tumor–draining lymph nodes of mice treated as in (A). Data shown represent two independent experiments with a total of 4-7 mice per group. (\**P* < 0.05, \*\*\**P* < 0.001, \*\*\*\**P* < 0.0001, two-way ANOVA with Holm-Sidak post hoc test). (**E**) Eight-week-old CD45.1 mice were irradiated, then reconstituted with bone marrow (BM) from Bcl6^fl/fl^CD4-cre BL/6 CD45.2 mice. Seven weeks post reconstitution, chimeric mice and age-matched C57/BL6 wild-type controls were implanted subcutaneously into each hind flank with MB49 cells (2 x 10^5^). When tumor volumes reached ∼100 mm^3^ (day 9), chimeric and control mice were treated with a single intratumoral injection of 1 x 10^6^ CFU of EcN^hCXCL13^. Intraperitoneal injections of PD-1 blockade were performed on day 12, 15, and 18. Tumor growth was monitored until day 41. (**F**) Subcutaneous MB49 tumor growth curves from Bcl6^fl/fl^CD4-cre BL/6 chimeric mice or age-matched C57/BL6 wild-type control mice (*n*=3-4 per group) that received intratumoral injections of EcN^hCXCL13^ and intraperitoneal PD-1 blockade as in (E). Data represents two independent experiments (\*\*\*\**P* < 0.0001, two-way ANOVA with Holm-Sidak post hoc test).

The co-stimulatory molecule ICOS and the inhibitory receptor PD-1 play a central role in establishing the Tfh program and controlling the homing and positioning of CD4^+^ T cells in the B cell follicle during the GC response (*14, 28*). Our analysis revealed that ICOS and PD-1 co-expression significantly increased in CD4^+^ helper T cells after intravesical delivery of EcN^hCXCL13^ as compared with EcN^−^, BCG, or PBS **(Fig.3C, and Fig.3D)**. However, we did not observe any upregulation of CXCR5, similar to previous observations in Tfh cells from human cancer–associated tissues **(fig.S3A)** (*29, 30*). Of note, the median survival of mice treated with EcN^hCXCL13^ alone was 53 days, as compared with 39 days and 37 days in mice treated with EcN^−^ or BCG, respectively **(fig.S3, B and C)**. These observations confirm that EcN^hCXCL13^ fosters the activation of CD4^+^ T helper cells with a pre-Tfh phenotype but has limited therapeutic effect when used as a single therapeutic agent.

Next, we hypothesized that activated CD4^+^ cells expressing high levels of PD-1 molecules may respond to PD-1 blockade. To test this in vivo, we assessed the frequency of Tfh cells co-expressing PD-1 and the Tfh-defining transcription factor Bcl6 following intravesical delivery of EcN^hCXCL13^ combined with PD-1 blockade. In this setting, we observed a significant increase in the frequencies of bladder tumor–draining lymph node Tfh cells (Bcl6^+^PD-1^+^) upon treatment with EcN^hCXCL13^ plus PD-1 blockade, as compared to mice treated with EcN^−^ or PBS plus PD-1 blockade, suggesting a dynamic effect of PD-1 blockade on Tfh differentiation **(fig.S3, D and E)**. Of note, our analyses also revealed a significant increase in the frequencies and total numbers of CD4^+^CD62L^+^CD44^+^ bladder tumor–draining lymph node central memory T cells in mice treated with combination EcN^hCXCL13^ plus PD-1 blockade therapy, which further highlights a potential role of memory T cells induced by our engineered probiotic therapy in the dynamics of the GC reaction **(fig.S3F)**.

To further investigate the role of Tfh cells in driving the therapeutic efficacy of EcN^hCXCL13^ plus PD-1 blockade, we generated animals deficient in Tfh generation by transplanting bone marrow from *Bcl6^fl/fl^*CD4-cre mice into lethally irradiated C57BL/6 recipient animals. Following reconstitution, MB49 cells were subcutaneously implanted into the hind flanks of Tfh-deficient bone marrow chimeras or age-matched wild-type control animals. At 9 days post-engraftment, mice received a single intratumoral injection of EcN^hCXCL13^, followed by intraperitoneal injections of anti-PD-1 antibody **(Fig.3E)**. As in the orthotopic setting, treatment with EcN^hCXCL13^ plus PD-1 blockade mediated potent antitumor control in wild-type; however, in bone marrow chimeric mice lacking Tfh cells (*Bcl6^fl/fl^*CD4-cre), the therapeutic benefit of EcN^hCXCL13^ plus PD-1 blockade combination therapy was abolished **(Fig.3F)**. Altogether, these findings demonstrate that EcN^hCXCL13^ plus PD-1 blockade induces the differentiation of CD4^+^ T helper cells into Tfh cells, which ultimately mediate the antitumor efficacy afforded by this combination therapy.

### EcN^hCXCL13^ plus PD-1 blockade boost the production of antitumor antibodies

We next aimed to understand the consequences of Tfh differentiation and expansion on the GC reaction in bladder tumor–draining lymph nodes. To do this, we assessed the immune phenotype of GC B cells by flow cytometry following treatment of MB49 bladder tumor–bearing mice with intravesical EcN^hCXCL13^, EcN^−^, or PBS, plus PD-1 blockade **(Fig.4A)**. As compared to mice receiving EcN^−^ or PBS plus PD-1 blockade, upon treatment with EcN^hCXCL13^ plus PD-1 blockade, we observed a marked increase in the frequencies of B220^+^CD19^+^GL7^+^CD95^+^ GC B cells, along with increased frequencies of proliferating GC B cells (Ki-67^+^Bcl6^+^) **(Fig.4, B, C, and D)**. These data demonstrate that EcN^hCXCL13^ plus PD-1 blockade enhances the GC reaction in bladder tumor–draining lymph nodes.

**Figure 4.**
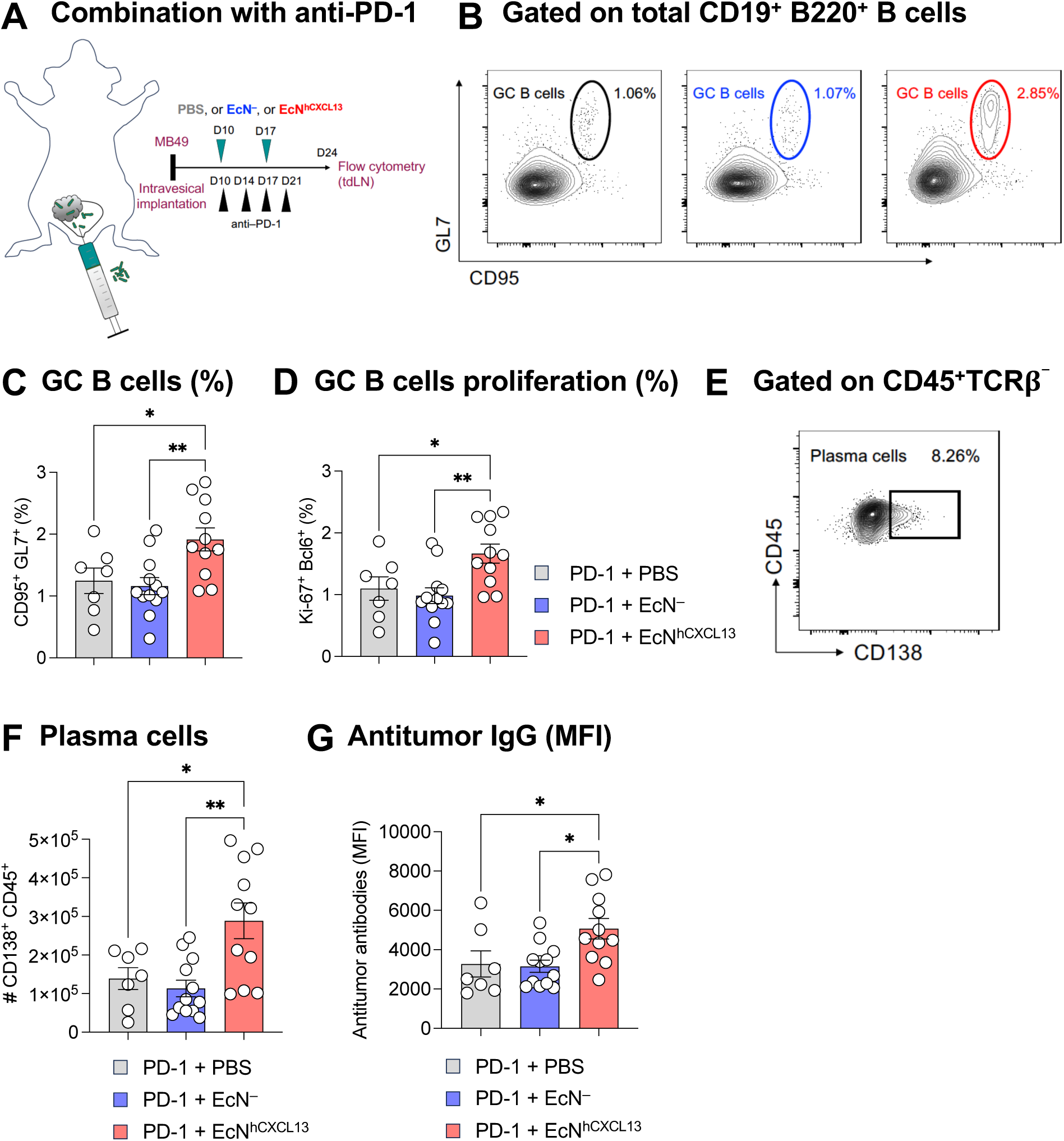
EcN^hCXCL13^ plus PD-1 blockade boosts the production of IgG antitumor antibodies. (**A**) MB49 cells (2 x 10^5^) were implanted in the bladder of wild-type C57BL/6 female mice. Intravesical delivery of either 3 x 10^6^ CFU of EcN_slic_^hCXCL13^, EcN^−^ control (empty vector), or PBS was performed on day 10 and 17. Intraperitoneal injections of PD-1 blockade were done on day 10, 14, 17, and 21. Bladder tumor–draining lymph nodes were harvested and dissociated at day 24 for immune phenotyping by flow cytometry. (**B**) Quantification of CD95^+^GL7^+^ germinal center B cells among total B cells in bladder tumor–draining lymph nodes. Data shown represent two independent experiments with a total of 3-5 mice per group. (\*\*\*\**P* < 0.0001, two-way ANOVA with Holm-Sidak post hoc test). (**C**) Representative dot plots showing Bcl6^+^Ki-67^+^ proliferative germinal center B cells among total B cells in draining lymph nodes from bladder-tumor bearing mice. (**D**) Quantification of Bcl6^+^Ki-67^+^ proliferative germinal center B cells among total B cells in bladder tumor–draining lymph nodes. Data shown represent two independent experiments with a total of 3-5 mice per group. (\*\*\*\**P* < 0.0001, two-way ANOVA with Holm-Sidak post hoc test). (**E**) Representative dot plots showing activated CD138^+^ plasma cells among CD45^+^TCRβ^-^ cells. (**F**) Quantification of the total number of plasma cells in bladder tumor–draining lymph nodes in the treated groups. Data shown represent two independent experiments with a total of 3-5 mice per group. (\*\*\*\**P* < 0.0001, two-way ANOVA with Holm-Sidak post hoc test). (**G**) MB49 cells were incubated with the serum of mice from each treatment group. The Ig binding affinity of IgG antibodies was quantified by mean fluorescence intensity.

To evaluate GC B cell differentiation into antibody-secreting cells, we assessed the frequency of CD138^+^ plasma cells in bladder tumor–draining lymph nodes. Analysis of CD45^+^ immune cells by flow cytometry showed a ∼3-fold increase of plasma cells upon EcN^hCXCL13^ plus PD-1 blockade therapy, as compared to EcN^−^ or PBS plus PD-1 blockade **(Fig.4, E and F**). These data suggest that actively proliferating GC B cells expand in response to EcN^hCXCL13^ plus PD-1 blockade and further differentiate into plasma cells. To investigate the potential production of tumor-specific antibodies by plasma cells, endpoint sera from each treatment group of MB49 bladder tumor–bearing mice were incubated in vitro with cultured MB49 cells. Whereas the total level of circulating IgG antibodies was similar in each treatment group, as compared to mice treated with EcN^−^ or PBS plus PD-1 blockade, a significant increase in the specificity and/or binding affinity of serum antibodies for MB49 cells was observed for sera isolated from mice treated with EcN^hCXCL13^ plus PD-1 blockade **(Fig.4G, fig.S4A)**. Of note, circulating IgM and IgA antibodies did not demonstrate any detectable binding to MB49 bladder cancer cells **(fig.S4B)**. Together, these observations demonstrate that EcN^hCXCL13^ plus PD-1 blockade promotes the GC reaction and plasma cell differentiation, resulting in the production of tumor-specific IgG antibodies in an immunocompetent model of bladder cancer.

## Discussion

Collectively, we have demonstrated the safety and efficacy of an approach which combines local delivery of engineered probiotics and systemic PD-1 blockade to promote antitumor immunity in orthotopic models of bladder cancer. This is notable, given that the MB49 model is an aggressive, fast-growing model of bladder cancer, and UPPL1541 tumors have been characterized as immunologically cold and resistant to PD-1 blockade (*24, 26*). We have shown that CXCL13-expressing *E. coli* colonize bladder tumors upon intravesical delivery to promote the GC response and the production of tumor-specific antibodies in tumor-draining lymph nodes upon combination with PD-1 blockade.

Innate detection of live *E. coli*, but not heat-killed bacteria, constitutes a physiological trigger for instruction of Tfh differentiation, an evolutionary conserved response in mice and humans (*31–33*). GCs are strictly dependent on the presence of T helper cells, in that they fail to form or be properly maintained in mice and humans deficient in CD4^+^ T cells or key components of T helper cell help (*4*). Here, by harnessing the tumor-restricted growth of non-pathogenic *E. coli* Nissle 1917 (EcN), we observed a potent GC reaction that amplified the production of antitumor antibodies following treatment with EcN^hCXCL13^ plus PD-1 blockade, while avoiding bacterial colonization of non-tumoral bladder tissue. Mechanistically, the CXCL13 gradient and the potential domination of ICOS signaling over PD-1 signaling in PD-1 blockade–treated mice may facilitate the appropriate migration of T helper cells to engage T cell–B cell contacts (*28, 34*). Our findings are aligned with this principle and provoke further interest in promoting (i) expansion of tumor-specific T follicular helper (Tfh) cells, (ii) homing of Tfh cells to the B cell follicles to enhance cognate B cell help through T–B interactions, and (iii) induction of a humoral immune response against tumor-specific antigens. In addition, the sharp increase in memory CD4^+^ T cells during the GC response suggests a critical role for memory T cells in the context of GC formation upon treatment with EcN^hCXCL13^ plus PD-1 blockade (*35*). These findings highlight the potential of engineering bacteria to promote long-term protective immunity as evidenced by our findings showing tumor rejection upon rechallenge in mice treated with EcN^hCXCL13^ plus PD-1 blockade.

In the context of cancer immunotherapy, the role of humoral immune responses in secondary lymphoid organs remains poorly investigated, mostly because lymphoid tissue is inaccessible in human cohorts (*18*). Recent studies have identified tumor-infiltrating B cells and organized T and B cell immune aggregates called tertiary lymphoid structures (TLS) as strong predictors of response to ICB in multiple solid tumors including bladder cancer (*36–42*). Indeed, mature TLS with GC-containing B cell follicles are required to enhance ICB responsiveness, most likely due to the production of high affinity antitumor antibodies (*39, 43, 44*). However, most solid tumors lack TLS or form immature TLS, leading to incomplete B cell differentiation and a shift toward a non-canonical GC pathway, with extra-follicular responses and low-affinity antibody-secreting cells (*45*). Alternative therapeutic strategies are evaluating the induction of TLS in the TME to enhance antitumor immunity independently of ICB immunotherapy (*46–48*).

Overall, our data support the notion that bacterial colonization of bladder tumors by EcN^hCXCL13^ combined with PD-1 blockade enhance the production of tumor-specific IgG antibodies by boosting the GC reaction in tumor-draining lymph nodes. Understanding the dynamics of Tfh differentiation and migration to the sites of B cell antibody production of tumor-specific antibodies will be critical for the development of potent bacterial cancer therapies and the design of combinatorial therapeutic approaches with ICB.

## Materials and Methods

### Study design

The objective of the study was to develop a probiotic intravesical delivery platform for intratumoral release of the human chemokine CXCL13 designed to induce GC B cell reaction upon colonization of bladder tumors. The experiments were designed to first assess whether the bacteria could locally produce and release a functional chemokine in bladder tumors and draining lymph nodes and then assess the efficacy of the probiotic platform in vivo in orthotopic mouse models of bladder cancer with and without combination with PD-1 blockade. The in vivo antitumor efficacy of the probiotic platform was assessed in MB49 and UPPL1541 syngeneic and orthotopic bladder tumors. Studies in mice were carried out to address the mechanisms through which these therapies function in combination with PD-1 blockade. All mice were randomized before treatment, and tumor volumes were measured by ultrasound imaging of the bladder post tumor engraftment. Groups were assigned with matching tumor burden between experimental conditions. Colonization efficiency, immune profiling by flow cytometry, and immunohistochemistry were used to characterize the system in vivo. Mouse body weight was monitored as a proxy for mouse health. Statistical analysis and sample sizes were determined from previous studies (*19–22*). Further details on sample size and replications (technical or biological) are provided in figure legends.

### Mouse strains

All experiments were performed in compliance with institutional guidelines and were approved by the Columbia University Institutional Animal Care and Use Committee (protocol AC-AABD8554). Seven-week-old wild-type C57BL/6NJ mice were purchased from the Jackson Laboratory, allowed to acclimate for a week, and then injected with cancer cells. Mice were randomized into treatment groups. All mice used were 8 to 12-week-old at time of experiment. Mice were housed at the Herbert Irving Comprehensive Cancer Center or at the Hammer Health Sciences Building under specific pathogen–free conditions.

### Cell lines

The MB49 mouse bladder cancer cell line was purchased at EMD Millipore Sigma (Catalogue number SCC148) (*25*). The UPPL1541 mouse bladder cancer cell line was a gift from W. Kim (University of North Carolina at Chapel Hill) (*26*). The SU-DHL-4 human B cell lymphoma cell line was purchased from The American Type Culture Collection (ATCC CRL-2957) (*23*). All cell lines tested negative for mycoplasma contamination. The MB49 and UPPL1541 cell lines were cultured inside tissue culture– treated flasks with Dulbecco’s modified Eagle medium (DMEM) with 10% fetal calf serum (HyClone) and 1% penicillin-streptomycin (100 U/ml; Gibco), nonessential amino acids, Gluta-MAX, Hepes, sodium pyruvate, and beta-mercaptoethanol. The SU-DHL-4 cell line was cultured in complete RPMI supplemented with 20% FCS and 10% DMSO. Cultures were maintained within a humidified 5% O2/CO2 atmosphere at 37°C inside an incubator. Cells were split twice per week, and cell viability was measured using trypan blue staining.

### Bacteria generation and preparation for intravesical delivery

The mature bioactive region of the human chemokine CXCL13 (hCXCL13) (Val^23^–Arg^94^, UniProt accession number Q853X90) or the mouse chemokine (mCXCL13) (Ile22-Ala-109, UniProt accession number Q3U1E8) were cloned into plasmid p246 via Gibson Assembly. CXCL13-expressing vectors were transformed into electrocompetent EcN-SLIC strains and cultured in LB media with 50 μg/ml kanamycin with 0.2% glucose, in a 37°C shaking incubator. For therapeutic preparation, EcN^hCXCL13^ and EcN^−^ (empty vector) strains were grown overnight in LB media containing appropriate antibiotics and 0.2% glucose. The overnight culture was sub-cultured at a 1:100 dilution in 50 mL of fresh media with antibiotics and glucose and grown to an OD600 of ∼ 0.05, preventing bacteria from reaching quorum. Cells were centrifuged at 3000 rcf and washed 3 times with sterile ice-cold PBS. EcN^hCXCL13^ and EcN^−^ strains were then diluted to a final concentration of 3×10^7^ CFU/mL in cold PBS, and 100 μL of each strain (3 x 10^6^ CFU) was then injected intravesically. Bacillus Calmette-Guerin (BCG, TICE strain, Merck) was obtained by resuspension in PBS of the lyophilized content of clinically available MSD vials. BCG was delivered into the bladder in a 100 μl volume of phosphate-buffered saline (PBS) containing 3 x 10^6^ CFU of bacteria.

### Intravesical tumor implantation

MB49 and UPPL1541 cells were detached from tissue culture plastic using 0.05% trypsin-EDTA (Gibco) for 1 to 3 min at 37°C. Cells were then washed using PBS, and cell viability and counting was assessed using trypan blue staining. Cells were resuspended in PBS at 2 x 10^6^ cells/mL for MB49 or 15 x 10^6^ cells/mL for UPPL1541. Eight-to 12-week-old female mice (The Jackson Laboratory) were placed under anesthesia in an isoflurane chamber. A 24-gauge catheter (Terumo) was inserted into the bladder through the urethra. Next, 100 μl of poly-L-lysine (Sigma) was injected in the bladder through the catheter, the catheter was capped using an injection plug (Terumo), and the mice were kept under anesthesia for 30 min. The mouse was removed from the isoflurane chamber, the bladder was manually emptied, and the catheter was removed. The catheter was then flushed with a solution containing 2 x 10^5^ MB49 cells/mL in PBS or 15 x 10^6^ cells/mL for UPPL1541 in PBS. Next the catheter was reinserted and 100 μl of the MB49 solution (∼250,000 cells per mouse) or UPPL1541 solution (∼1,500,000 cells per mouse) was injected into the bladder. The mice were kept under anesthesia for 2 hours and were allowed to recover from anesthesia. Mice were then monitored regularly after implantation for signs of hematuria and tumor growth. To generate models of advanced bladder cancer, (I) MB49 cells were grafted 10 days before starting treatment; (II) UPPL1541 cells were implanted in the lamina propria by ultrasound-guided injection at 5 x 10^6^ cells suspended in 500 μL sterile PBS using 30G needle with syringe as previously described (*27*). Ultrasound imaging of the bladder was used to measure the tumor volume pre- and post-treatment. Groups were assigned with matching pre-treatment tumor volume between experimental conditions. Mice were weighed two times weekly and euthanized when the humane endpoint of 20% weight loss was reached, or if they displayed signs of distress, such as dull fur or apathy.

### Intravesical treatment

Mice were anesthetized, and 4-gauge plastic catheters were used to instill bacteria into the bladder of mice. A total volume of 100 μL of 3 x 10^6^ CFU of EcN^hCXCL13^, EcN^−^, or BCG was injected into the bladder. PBS was used as a control. The catheter was capped using an injection plug (Terumo). The mice were kept under anesthesia for 2 h. At the end of this time, catheters were removed, and the mice were allowed to recover from anesthesia. Dosing schemes are indicated in each figure and figure legends.

### Subcutaneous tumor implantation

To assess immune memory, surviving mice were subcutaneously rechallenged with MB49 cells after MB49 orthotopic implantation. All surviving mice were free of bladder tumors. Mice were injected on both flanks with 100 µL at 2 x 10^6^ cells/mL of MB49 cells (∼500,000 cells per mouse). Caliper measurements were used to track tumor volume every 3-4 days. Tumor volume was calculated by measuring the length, width and height of each tumor using calipers, where V=(length × width × height x π/6). Mice were euthanized when tumor volumes reached 1,000 mm^3^.

### Intratumoral treatment

For intratumoral bacterial injection, a single dose of EcN^hCXCL13^ strain (1 x 10^6^ CFU) in 40 μL of PBS was injected. Bacteria were injected on day 9 in subcutaneously implanted MB49 tumors in both hind flanks.

### Generation of bone marrow chimeric mice

To deplete T follicular helper cells (Tfh) in mice, we generated full bone marrow (BM) chimeras by reconstituting 8-week-old male C57BL/6 mice with BM from *Bcl6^fl/fl^*CD4-cre male C57BL/6 mice provided by Shane Crotty’s lab (Center for Infectious Disease and Vaccine Research, La Jolla Institute for Immunology, La Jolla, CA). To achieve complete myeloablation, 8-week-old wild-type C57BL/6NJ male mice were exposed to whole body irradiation (2 doses of 5.25 Gy per mouse, 4 h apart). After at least 4 h from the last irradiation, mice were reconstituted with 5 × 10^6^ whole BM cells isolated from donors and injected intravenously by retro-orbital injection. Mice were treated with antibiotics in the drinking water for 4 weeks after reconstitution. Mice were placed in standard cages and allowed to reconstitute for 4 more weeks total before subcutaneous tumor implantation.

### Ultrasound imaging

Mice were anesthetized using inhalation isoflurane, and hair was removed from their lower abdomen by using chemical hair removal cream. Ultrasound gel was applied to the abdomen, and mouse bladders were imaged using the VEVO 3100 Ultrasound Imaging System (FUJIFILM VisualSonics, Toronto, Canada) located within the mouse barrier in the Herbert Irving Cancer Center Small Animal Imaging facility. Tumor volume was calculated via 3D reconstruction program (Vevo LAB).

### Chemotaxis assay

To characterize the chemotaxis effect of hCXCL13-derived bacteria on mouse lymphocytes, splenocytes were isolated from wild-type adult C57BL/6 mouse spleen using mechanical dissociation in wash buffer (RPMI 1640 supplemented with 10% FBS, HEPES, Glutamax, Pen/Strep). Cells were filtered through 100um cell strainers and resuspended in wash media at a concentration of 1 x 10^6^ cells per mL. Overnight cultures of each bacterial strain (without SLIC) were grown in LB with appropriate antibiotics and then subcultured at a 1:100 dilution in a shaking incubator for 60 min at 37 °C. Bacteria were washed twice in serum-free complete RPMI, matched at optical density at 600 nm (OD600), and lysed via sonication in serum-free complete RPMI. Lysates were centrifuged to remove debris (20817g for 10 min at 4°C), and 235 μl of the supernatant was entered into the lower chamber of a Corning HTS transwell plate (well area=0.143 cm2; pore size=5 μm). Mouse lymphocytes (75 μl of the preparation described above) were added to the upper chamber, and the plate was incubated for 3 hours in a humidified 37°C 5% CO2 incubator. The bottom chamber was then harvested and washed in complete media. Then, samples were acquired on a BD Fortessa for 60 s. Cell counts were normalized to the number of cells entered into the assay. To characterize the chemotaxis effect of hCXCL13-derived bacteria on human B cells, SU-DHL4 cells were harvested from cell culture and used as mentioned above.

### Enzyme-linked immunosorbent assay

For in vitro quantification of human CXCL13, overnight cultures of EcN^−^ and EcN^hCXCL13^ were grown in LB agar with or without kanamycin, and then subcultured (1:100 dilution) for 60 minutes in a shaking incubator at 37°C. Bacteria were harvested, concentration matched at optical density at 600 nm (OD600), and resuspended in 2 mL LB, followed by continued culturing in a shaking incubator for 60 min at 37°C in presence of AHL for SLC-induced lysis(*20*). Cultures were then centrifuged (3000 g for 10 min at 4°C), and supernatants were entered into a human CXCL13 enzyme-linked immunosorbent assay (ELISA) (R&D Systems™ Human CXCL13/BLC/BCA-1 DuoSet ELISA, catalog number DY801), performed as per the manufacturer’s protocol. To quantify mouse IgG after combined therapy with PD-1 blockade and intravesical EcN^hCXCL13^, EcN^−^, or PBS, the serum levels of total circulating IgG antibodies were detected and quantified using the Mouse IgG (Total) uncoated ELISA kit (Catalog number 88-50400-22), performed as per the manufacturer’s protocol.

### Immunophenotyping by flow cytometry

#### Bladder processing

Bladders were dissected, cut into small pieces, and digested in wash media (RPMI 1640 supplemented with 5% FCS, HEPES, Glutamax, Pen/Strep) with 1 mg/mL collagenase A and 0.5 µg/mL DNAse I in a shaking incubator at 37°C for 45 minutes. Digestion was stopped by adding FACS buffer (PBS supplemented with 1% bovine serum albumin, 2.5 mM EDTA, and 0.1% NaN_3_). Single-cell suspensions were washed and resuspended in ice cold FACS buffer prior to staining.

#### Bladder-draining lymph nodes processing

Lymph nodes were homogenized with a syringe plunger and filtered through a 70 mm cell strainer on ice. Cells from all tissues were resuspended in ice cold FACS buffer and immediately used for counting or staining. Live/dead staining was performed via Ghost Dye Red 780 labeling (Tonbo Biosciences), as per the manufacturer’s protocol. All staining steps were carried out at 4°C in FACS buffer (PBS supplemented with 1% bovine serum albumin, 2.5 mM EDTA, and 0.1% NaN_3_).

Single cell suspensions were Fc blocked with anti-CD16/CD32 monoclonal antibody prior to staining. Cells were then stained for flow cytometry, with intracellular staining performed using the Tonbo Foxp3 /Transcription Factor Staining Buffer Kit as per the manufacturer’s protocol. Antibodies used included anti-CD45 (clone 30-F11, BD Biosciences), TCRβ (clone H57-597, BD Biosciences), B220 (clone RA3-6B2, BD Biosciences), CD19 (clone 1D3, Tonbo), GL7 (clone GL7, BD Biosciences), CD95 (clone JO2, BD Biosciences), CD138 (Clone: 281 2, BD Biosciences), CD4 (clone RM4-5, BD Biosciences), CD8 (clone 53-6.7, BioLegend), Foxp3 (clone FJK-16s, eBioscience), CD44 (clone IM7, BioLegend), CD62L (clone MEL-14, Tonbo), Bcl6 (clone K112 91, BD Biosciences), PD-1 (clone 29F.1A12, BioLegend), CXCR5 (clone L138D7, BioLegend), Ki-67 (SolA15, ThermoFischer), IgG (clone Poly4053, BioLegend), IgM (clone R6 60.2, BD Biosciences), IgA (clone C10-3, BD Biosciences). Total cell counts were determined by the addition of counting beads (Invitrogen) to a known volume of sample after staining, just before cytometer acquisition. All samples were acquired on a BD LSR-II or LSR-Fortessa using DIVA software, and data were analyzed in FlowJo (Tree Star) software.

### Immunostaining

Bladders were fixed overnight in 4% paraformaldehyde at 4°C and embedded in paraffin. Serial sections of 4 mm were generated and mounted on adhesive microscope slides. For immunofluorescence staining, paraffin sections were deparaffinized using HistoClear and rehydrated through a series of Ethanol and 1 X PBS washes. Antigen retrieval was performed by boiling slides for 15 min in pH 9 buffer or 30 min in pH 6 buffer. Primary antibodies in 1% heat inactivated horse serum and 1% bovine serum albumin were incubated overnight at 4°C. The next day, slides were washed with PBS–Triton 0.5% three times for 10 min each and secondary antibodies were applied for 90 minutes at room temperature. DAPI (4’6-diamidino-2-phenylindole) was applied as part of the secondary antibody cocktail for nuclear staining. Slides were sealed with coverslips using DAKO mounting gel and stored at 4°C until analysis.

### Fluorescent microscopy

Zeiss Axiovert 200M microscope with Zeiss apotome were used to collect immunofluorescent images.

### Quantification of bacterial viable cell numbers

Bladders were weighed and homogenized in 1 mL of sterile PBS at 4°C using a handheld rotor-stator tissue homogenizer (TissueRuptor II, Qiagen). Homogenates were serially diluted, plated on LB agar plates and incubated overnight at 37°C. For plasmid retention analysis, tumor homogenates were plated on LB-agar plates containing kanamycin. Colonies were counted and computed as CFU/mg of tissue.

### Detection of antitumor antibodies

Serum processing. Whole blood was collected via terminal cardiac puncture, incubated at room temperature for 45 min, and centrifuged at 14000 x g (maximum speed) for 15 minutes, 4°C. Serum was recovered and stored at - 20°C for antibody binding assays. To detect antitumor antibodies after therapy, MB49 cells were incubated with sera diluted 1:50 in PBS for 30 min at room temperature, washed with FACS buffer, stained with fluorescently labelled antibodies to mouse IgG, IgA and IgM for 30 min at room temperature and analyzed by flow cytometry on a LSR-II flow cytometer. Antibody titres are represented as the MFI per antibody isotype. Prior to incubation, MB49 cells were washed three times using centrifugation and PBS and resuspended in 100 μL PBS buffer. Sera were normalized at 40 ng/mL IgG in PBS. A pre-absorption protocol was followed to remove autoantibodies as previously described (*49*). Briefly, the pre-absorption protocol was done by incubating pre-diluted sera with cells from freshly dissociated normal bladder urothelium from C57BL/6NJ wild-type mice for 45 min on ice. Cells were then centrifuged, the supernatant collected and then incubated with MB49 cells for 45 min on ice. Background was assessed using the antibody against MB49 cells incubated with PBS–no serum.

### Statistical analysis

Statistical analysis was performed in GraphPad Prism 9, using Student’s T test or one- or two-way analysis of variance (ANOVA) with post hoc testing as indicated. A *P* value of <0.05 was considered statistically significant. The specific statistical test, sample sizes, and *P* values for each figure are indicated in the figure legends.

## Acknowledgments

We thank Shane Crotty for providing bone marrow from Bcl6^fl/fl^CD4-cre BL/6 mice, William Kim for providing the UPPL1541 cell line, and James M. McKiernan for providing the BCG vials. We are thankful for insightful discussion with the members of the Arpaia and Danino labs. We thank the members of the Mendelsohn lab for providing technical support with the orthotopic UPPL1541 model. This research used the resources of the Oncology Precision Therapeutics and Imaging Core at the Herbert Irving Comprehensive Cancer Center, and the Histology Service at the Columbia University Irving Medical Center.

## Author contributions

M.R. and N.A. conceived and designed the project with input from T.D. M.R., N.C., D.L.M., F.L., T.M.S, K.D.L.A., and R.L.V. designed and/or performed in vivo and ex vivo experiments. All the authors analyzed the data. C.L.M. provided input on the orthotopic models of bladder cancer and immunofluorescence imaging. M.R. and N.A wrote the manuscript with critical input from all authors.

---

**Supplementary Figure 1.**
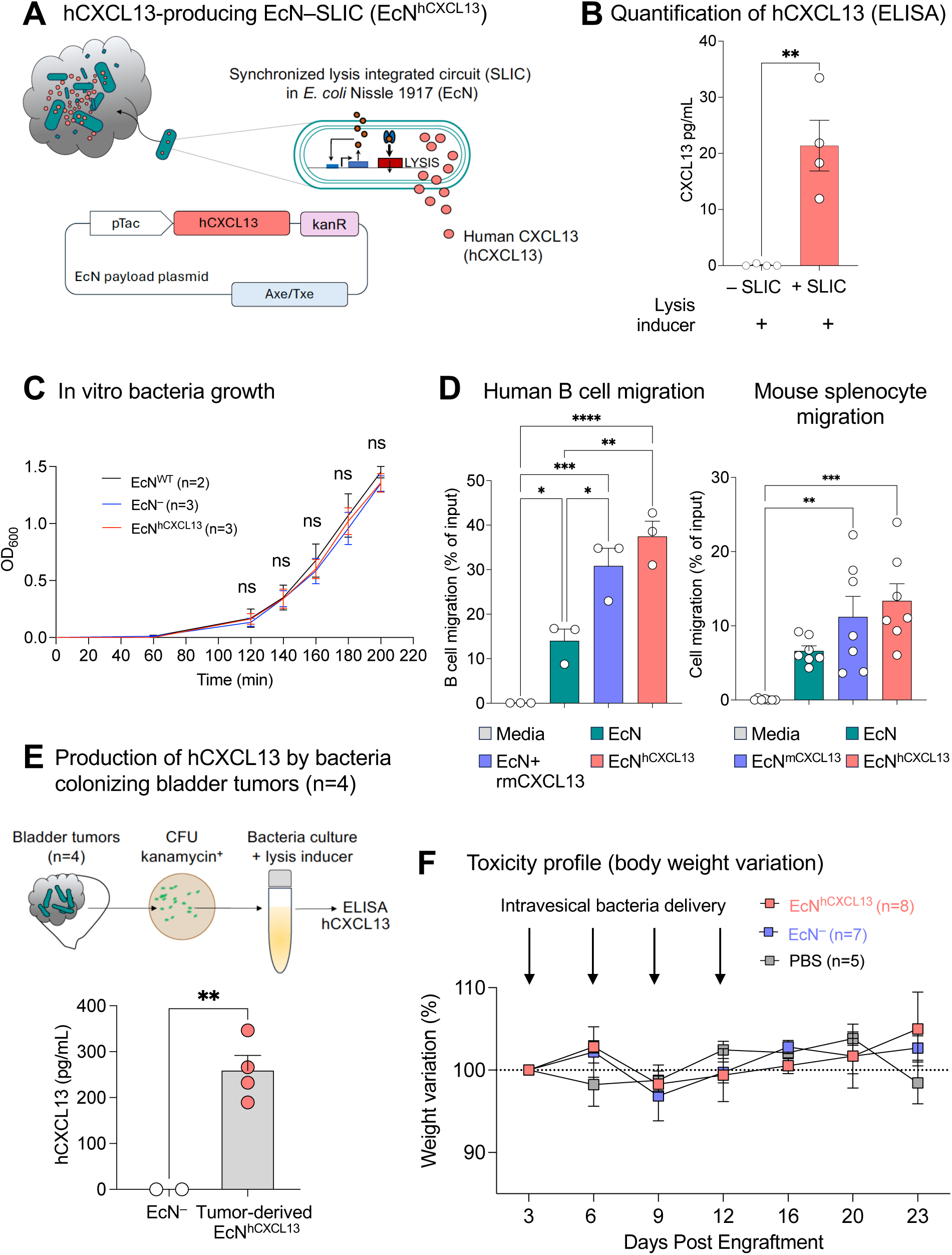
Engineered *E. coli* Nissle 1917 release a functional human chemokine CXCL13 for bladder cancer therapy. (**A**) Schematic demonstrating the release of the human chemokine CXCL13 in bladder tumors after tumor colonization by engineered probiotic *E.coli* Nissle 1917 upon intravesical delivery (left panel). PBS or MB49 cells (2 x 10^5^) were implanted in the bladder of C57/BL6 mice. Intravesical delivery of 3 x 10^6^ CFU of EcN^hCXCL13^ was performed on days 7, 14, and 21. On day 28, supernatant from freshly dissociated tumor-free (n=6) or tumor-bearing (n=5) bladders was plated on LB agar plates containing kanamycin. Colony-forming units (CFU) were counted after overnight culture at 37°C and computed as CFU per mg of bladder tissue. Bladder tumor–draining lymph nodes were dissociated for immune phenotyping by flow cytometry (right panel). (**B**) Quantification of the total number of CFU per bladder weight (mg) after overnight culture at 37°C on LB agar plate with kanamycin. Data shown represent two independent experiments with a total of 5-6 mice per group (**P < 0.01, two-tailed unpaired Mann Whitney test). (**C**) Representative dot plots showing total CD19^+^B220^+^ B cells in draining lymph nodes from bladder tumor-free (left panel) and bladder tumor-bearing (right panel) mice. (**D**) Flow cytometric quantification of total CD19^+^B220^+^ B cells among live CD45^+^ cells in draining lymph nodes from bladder-tumor-free and bladder-tumor bearing mice (*P < 0.05, two-tailed unpaired Student’s *t* test). (**E**) Flow cytometric quantification of CD95^+^GL7^+^ germinal center B cells among total B cells in draining lymph nodes from bladder tumor-free and bladder tumor-bearing mice (two-tailed unpaired Student’s *t* test). (**F**) Representative dot plots showing Bcl6^+^PD-1^+^ T follicular helper cells among TCRβ^+^CD4^+^FoxP3^-^cells in draining lymph nodes from bladder tumor-free (left panel) and bladder tumor-bearing (right panel) mice. (**G**) Flow cytometric quantification of Bcl6^+^PD-1^+^ T follicular helper cells among TCRβ^+^CD4^+^FoxP3^-^ cells in draining lymph nodes from bladder tumor-free and bladder tumor-bearing mice (**P < 0.01, two-tailed unpaired Student’s *t* test). (**H**) Mean fluorescence intensity (MFI) quantification of PD-1 molecules expressed at the surface of Bcl6^+^PD-1^+^ T follicular helper cells by flow cytometry in draining lymph nodes from bladder tumor-free and bladder tumor-bearing mice (**P < 0.01, two-tailed unpaired Student’s *t* test).

**Supplementary Figure 2.**
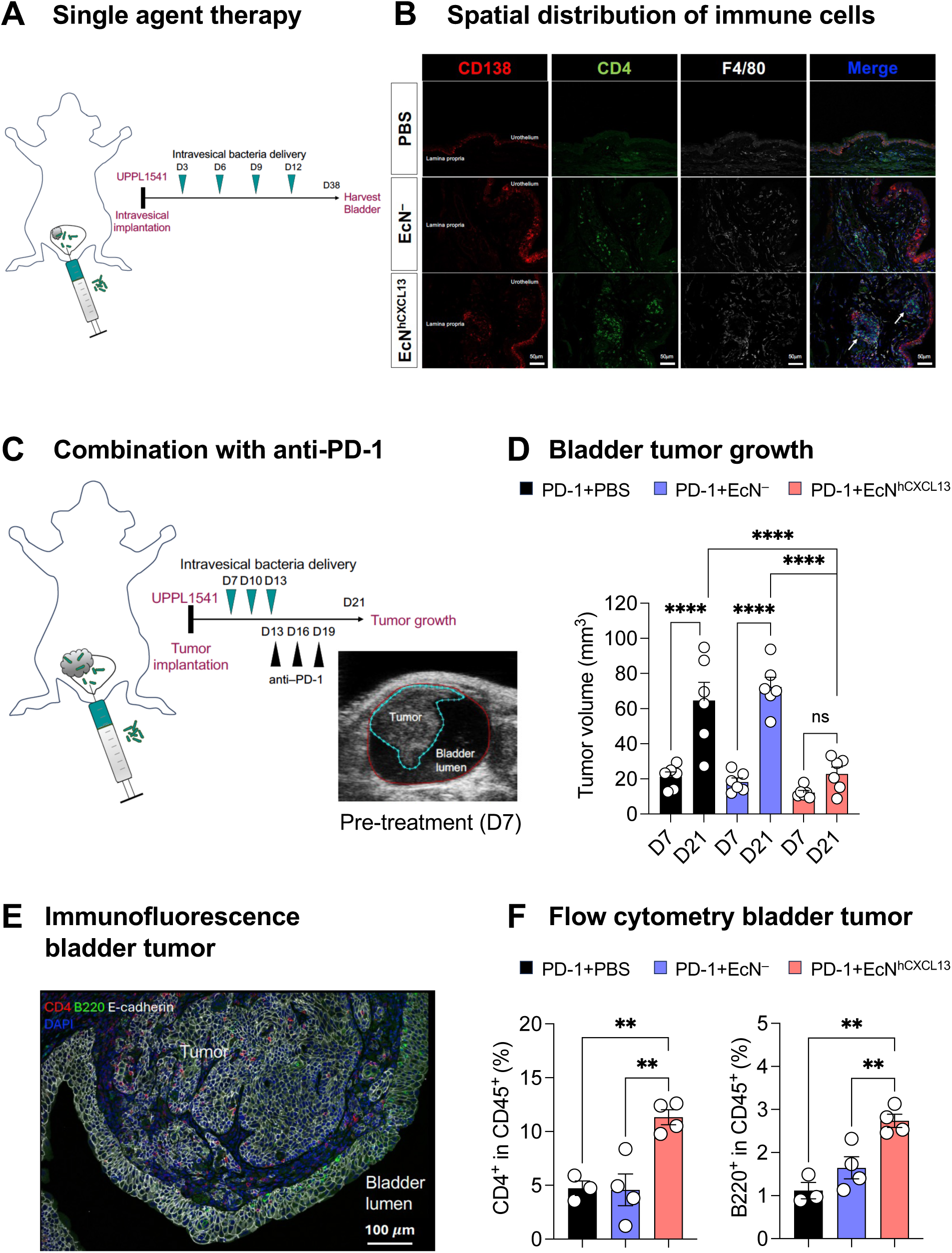
EcN^hCXCL13^ plus PD-1 blockade slow tumor progression and promote CD4^+^ T cells and B cells and infiltration in an orthotopic model of PD-1 resistant bladder cancer. (**A**) UPPL1541 cells (10 x 10^6^) were implanted in the bladder of wild-type C57BL/6 female mice. Intravesical delivery of either 3 x 10^6^ CFU of EcN^hCXCL13^, EcN^−^ control (empty vector), or PBS was performed on day 3, 6, 9 and 12. Bladders were harvested at day 38 for immunofluorescence imaging. (**B**) Representative images of whole bladder sections. Bladders were stained with monoclonal antibodies against CD138 (red), CD4 (green), F4/80 (white). Images display each channel respectively and merged signals; bar, 50μm. (**C**) UPPL1541 cells (5 x 10^5^) were implanted in the bladder submucosa of wild-type C57BL/6 female mice using an ultrasound-guided approach. Intravesical delivery of either 3 x 10^6^ CFU of EcN_slic_^hCXCL13^, EcN^−^ control (empty vector), or PBS was performed on day 7, 10, and 13. Intraperitoneal injections of PD-1 blockade were done on day 13, 16, and 19. On day 7 (pre-treatment) and day 21 (post-treatment), the volume of bladder tumors was monitored by ultrasound imaging. (**D**) Day 7 (pre-treatment) and day 21 (post-treatment) tumor volumes of wild-type C57BL/6 female mice are shown. Data represent two independent experiments with n=6 mice per group (****P < 0.0001, one-way ANOVA with Holm-Sidak post hoc test). (**E**) Bladder tumors were treated as in (**C**) and resected 48hrs after the last injection of PD-1 blockade for analysis by immunofluorescence and flow cytometry. Representative image of whole bladder section from a mouse treated with EcN^hCXCL13^ plus PD-1 blockade from one experiment with n=3-4 mice per group. Bladders were stained with monoclonal antibodies against CD4 (red), B220 (green), E-cadherin (white); bar, 100μm. (**F**) Frequency of CD45^+^TCRβ^+^CD4^+^ cells and CD45^+^TCRβ^-^B220^+^ cell populations from bladder tumors treated as in (**C**) are shown. Data represent one experiment with mean ± SEM of n=3-4 mice per group (**P < 0.01, one-way ANOVA with Holm-Sidak post hoc test).

**Supplementary Figure 3.**
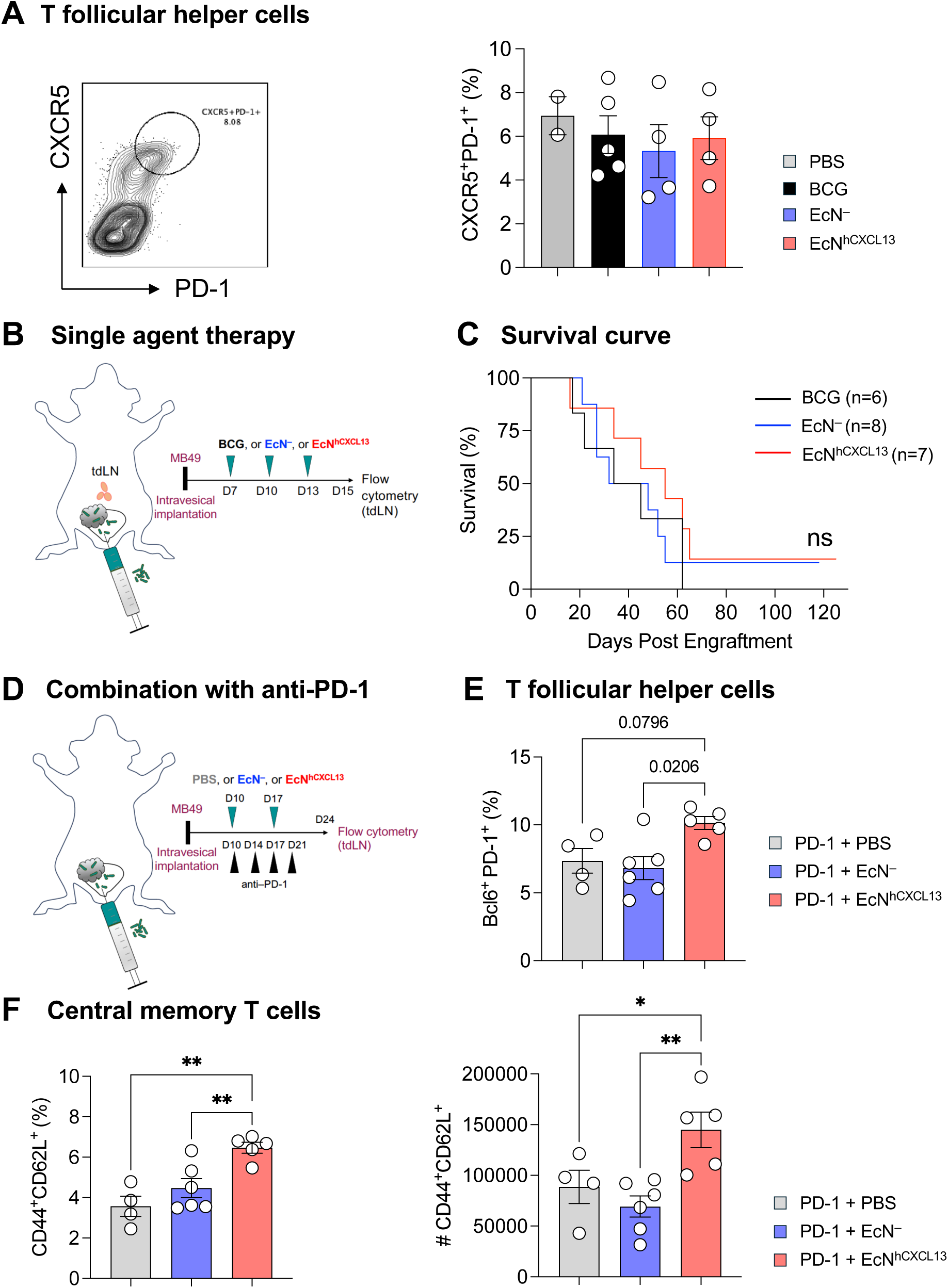
EcN^hCXCL13^ plus PD-1 blockade increase central memory T cells in an orthotopic model of advanced bladder cancer. (**A**) Representative flow plots (left panel) and frequency (right panel) of CXCR5^+^PD-1^+^ T follicular helper cells among TCRβCD4^+^FoxP3^-^ cells from bladder tumor–draining lymph nodes of mice treated as in (B) Bladder tumor–draining lymph nodes were harvested and dissociated for immune phenotyping 48 hours after the last intravesical delivery. (**B**). MB49 cells (2 x 10^5^) were implanted in the bladder of wild-type C57BL/6 mice. Intravesical delivery of either 3 x 10^6^ CFU of EcN^hCXCL13^, EcN^−^ control (empty vector), or BCG was performed at days 7, 10, and 13. Survival was monitored > day 100. (**C**) Survival curves are shown for each treatment group. Data represent two independent experiments with n=6-8 mice per group (log-rank test). (**D**) MB49 cells (2 x 10^5^) were implanted in the bladder of wild-type C57BL/6 female mice. At day 10, the volume of bladder tumors was monitored by ultrasound imaging. Intravesical delivery of either 3 x 10^6^ CFU of EcN^hCXCL13^, EcN^−^ control (empty vector), or PBS was performed at day 10, 17, 24, and 31. Intraperitoneal injections of PD-1 blockade were done on day 10, 14, 17, and 21. Bladder tumor–draining lymph nodes were harvested at day 24 for immune phenotyping by flow cytometry. (**E**) Quantification of Bcl6^+^PD-1^+^ among TCRβ^+^CD4^+^FoxP3^-^ cells in bladder tumor–draining lymph nodes. Data shown represent one experiment with a total of 4-6 mice per group. (\**P* < 0.05, two-way ANOVA with Holm-Sidak post hoc test). (**F**) Quantification of CD44^+^CD62L^+^ central memory T cell frequency among TCRβ^+^CD4^+^FoxP3^-^cells (left panel) and number (right panel) in bladder tumor–draining lymph nodes. Data represent one independent experiment with 4-6 mice per group. (\**P* < 0.05, \*\**P* < 0.01, two-way ANOVA with Holm-Sidak post hoc test).

**Supplementary Figure 4.**
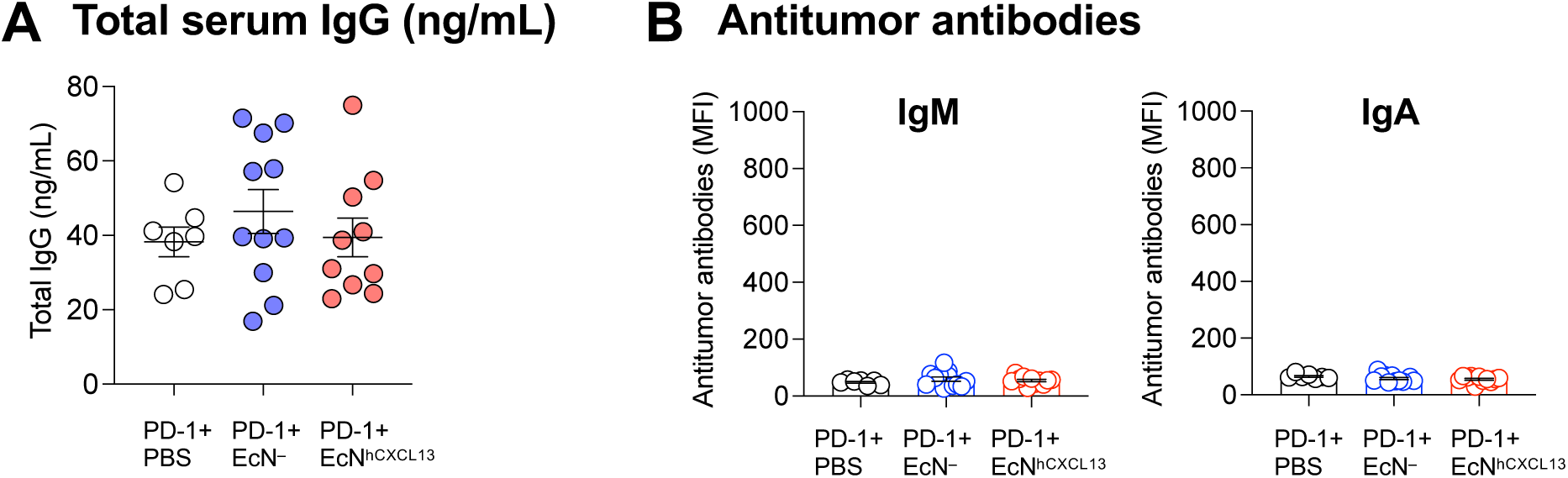
EcN^hCXCL13^ plus PD-1 blockade do not induce IgA and IgM antitumor antibodies. (**A**) MB49 cells (2 x 10^5^) were implanted in the bladder of wild-type C57BL/6 female mice. Intravesical delivery of either 3 x 10^6^ CFU of EcN_slic_^hCXCL13^, EcN^−^ control (empty vector), or PBS was performed on day 10 and 17. Intraperitoneal injections of PD-1 blockade were done on days 10, 14, 17, and 21. The total level of circulating IgG antibodies was quantified by ELISA. (**B**) MB49 cells were incubated with the serum of mice from (**A**) for each treatment group. The Ig binding affinity of IgM and IgA antibodies was quantified by mean fluorescence intensity.

